# Fine Specificity Epitope Analysis by HX-MS Identifies Contact Points on Ricin Toxin Recognized by Protective Monoclonal Antibodies

**DOI:** 10.1101/345561

**Authors:** Greta Van Slyke, Siva Krishna Angalakurthi, Ronald T. Toth, David J Vance, Yinghui Rong, Dylan Ehrbar, Yuqi Shi, C. Russell Middaugh, David B. Volkin, David D. Weis, Nicholas J. Mantis

**Affiliations:** Division of Infectious Disease, Wadsworth Center, New York State Department of Health, Albany, NY 12208; Department of Pharmaceutical Chemistry and Macromolecule and Vaccine Stabilization Center, University of Kansas, Lawrence, KS 66045; Department of Chemistry and Ralph Adams Institute for Bioanalytical Chemistry, University of Kansas, Lawrence, KS 66045

**Keywords:** toxin, antibody, epitope, biodefense

## Abstract

Ricin is a fast-acting protein toxin classified by the Centers for Disease Control and Prevention as a biothreat agent. In this report we describe five new mouse monoclonal antibodies (mAbs) directed against an immunodominant region, so-called epitope cluster II, on the surface of ricin’s ribosome-inactivating enzymatic subunit, RTA. The five mAbs were tested alongside four previously described cluster II-specific mAbs for their capacity to passively protect mice against 10 × LD_50_ ricin challenge by injection. Only three of the mAbs (LE4, PH12 and TB12) afforded protection over the seven-day study period. Neither binding affinity nor *in vitro* toxin-neutralizing activity could fully account for LE4, PH12 and TB12’s potent *in vivo* activity relative to the other six mAbs. However, epitope mapping studies by hydrogen exchange-mass spectrometry (HX-MS) revealed that LE4, PH12 and TB12 shared common contact points (*i.e.*, “strong” protection by HX-MS) on RTA that encompassed residues 154-164 and 62-69, which correspond to RTA α-helices D-E and β-strands d-e, respectively, located on the back side of RTA relative to the active site. The other six mAbs recognized overlapping epitopes on RTA but none shared the same HX-MS profile as LE4, PH12 and TB12. A high-density competition ELISA with a panel of ricin-specific single domain camelid antibodies (VHHs) indicated that even though LE4, PH12 and TB12 make contact with similar secondary motifs, they ultimately approach RTA different from angles. These results underscore how subtle differences in epitope specificity have significant impacts on the antibody functionality *in vivo* and have important implications in the design of immune-based countermeasures against ricin.

## Introduction

Ricin is at the top of the list of potential biothreat agents, according to a NATO Biomedical Advisory council (1). Ricin toxin is a product of the castor bean plant (*Ricinus communis*), which is cultivated worldwide for its oils used in industrial and cosmetic applications. The toxin itself is a ~65 kDa glycoprotein consisting of two subunits, RTA and RTB, joined by a single disulfide bond (2). RTA is an extraordinarily efficient RNA Nglycosidase (EC 3.2.2.22) that cleaves the sarcin-ricin loop (SRL) of 28S rRNA, resulting in ribosome inactivation (3, 4). RTB is a galactose/N-acetyl galactosamine (Gal/GalNAc)-specific lectin that facilitates RTA endocytosis and retrograde transport from the plasma membrane to the endoplasmic reticulum (ER) of mammalian cells. In the ER, RTA is liberated from RTB, partially unfolded, and then retro-translocated across the ER membrane into the cell cytoplasm, presumably via the Sec61p translocon (5). In rodents and non-human primates, the lethal dose 50 (LD_50_) of ricin ranges from 1 to 10 μg/kg by injection or inhalation (6).

RTA is the focus of current efforts to develop a countermeasure for ricin, including a subunit vaccine for use by first responders and military personnel (7-9). RTA, 267 amino acid residues in length, is a globular protein with a total of 10 β-strands (a-j) and seven α-helices (A-G) (2, 10). The active site constitutes a shallow cleft on one side of the molecule. There are four distinct immunodominant regions or epitope clusters on the surface of RTA, originally identified through competition ELISAs with four different toxin-neutralizing monoclonal antibodies (mAbs) (11, 12, 13). Cluster I is focused around RTA’s α-helix B (residues 94-107), a protruding element previously known to be a target of potent toxin-neutralizing antibodies (14, 15). Cluster II is defined by the mAb SyH7 and is located on the back side of RTA, relative to the active site pocket. Cluster III involves α-helices C and G on the front side of RTA, while cluster IV forms a diagonal sash from the front to back of the A subunit.

Prior to this report, cluster II consisted of overlapping epitopes defined by four mouse mAbs: SyH7, PA1, TB12 and PH12 (13). All four mAbs have *in vitro* toxin-neutralizing activities and have been shown to passively protect mice from 5 × LD50 ricin challenge by injection over a 72 h period (11, 12). However, recent epitope mapping studies using hydrogen exchange-mass spectrometry (HX-MS) have indicated that cluster II actually sectors into at least two distinct sub-clusters (13). SyH7 engages RTA residues 14-24 (corresponding to α-helix A) and residues 184-207 (corresponding to a loop between α-helices F-G). PA1 also engages residues 184-207, while PH12 and TB12 contact RTA residues 62-69 (corresponding to a loop between β-strands d-e) and residues 154-164 (corresponding to a loop between α-helices D-E). Furthermore, we identified a collection of RTA-specific single domain camelid antibodies (V_H_Hs) that compete with SyH7, PA1, TB12 and PH12 to varying degrees for binding to ricin (16). The majority of the cluster II V_H_Hs are devoid of toxin-neutralizing activity. Overall, these results indicate that cluster II is much more complex than originally anticipated, encompassing a large amount of surface area on RTA with numerous binding sites for neutralizing and non-neutralizing antibodies. Therefore, the goal of the current study was to interrogate cluster II with additional mAbs in an effort to better define which specific structural elements are most associated with *in vivo* protection.

## Results

To generate addition cluster II mAbs, we screened B cell hybridomas derived from BALB/c and Swiss Webster mice that had been hyperimmunized with sub-lethal amounts of ricin toxin. Ricin-specific mAbs were identified by direct ELISA, while cluster II-specific mAbs were classified initially by competition ELISA with SyH7 (data not shown). In total we identified four new cluster II mAbs: LE4, CH1, SWB1, and 6C4 (**Table 1**). A fifth mAb, WECH1, was resurrected from a previous hybridoma screen (11). In addition to competition with SyH7, we performed cross competition analysis among the five new mAbs themselves (**Figure 1**), which confirmed that WECH1, LE4, CH1, SWB1, and 6C4 recognize overlapping epitopes on RTA.

**Figure 1.**
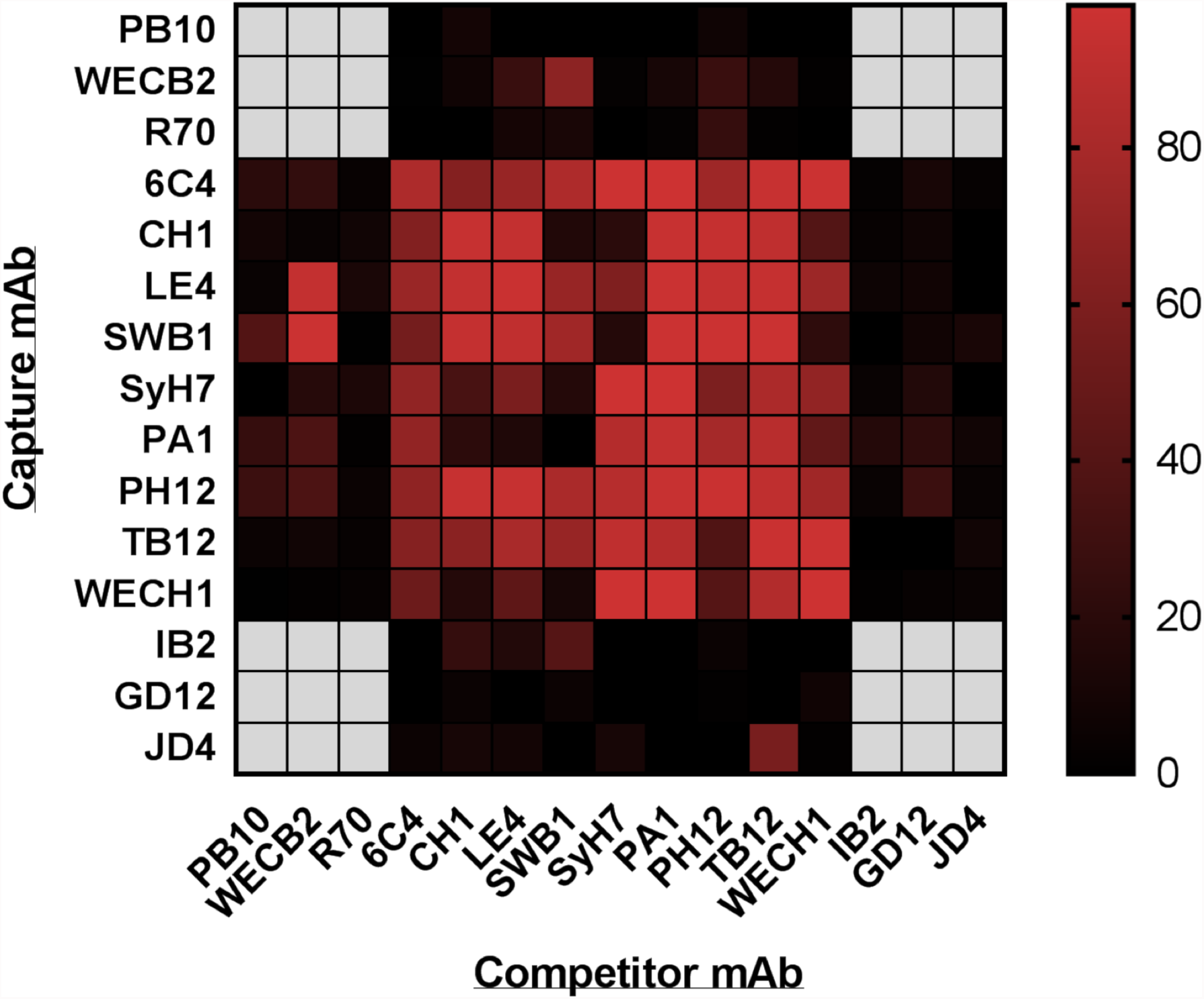
Binning by cross-competition ELISA. A heat map representation of a cross-competition ELISA with panel of RTA-specific mAbs, as described in the Materials and Methods. The mAbs listed on the vertical axis were coated onto microtiter plates and then assessed for the ability to capture soluble biotin-ricin in the presence of the indicated competitor mAb (horizontal axis). The percent (%) inhibition of biotin-ricin capture was calculated from the optical density (OD) values as follows: 1- value OD_450_ (biotin-ricin + competitor mAb)/ value OD_450_ (biotin-ricin without competitor mAb) × 100. The values were plotted as a heat map using Prism 7 (GraphPad). The scale bar on the right indicates percent inhibition from no competition (black) to complete competition (dark red). The heat map is presented as a means of visualizing the relative competition groups or clusters (I-IV) referred to in the body of the manuscript. PB10, WECB2 and R70 are in cluster I, IB2 in cluster III, and GD12 and JD4 in cluster IV. The remaining mAbs are in cluster II.

**Table 1.**
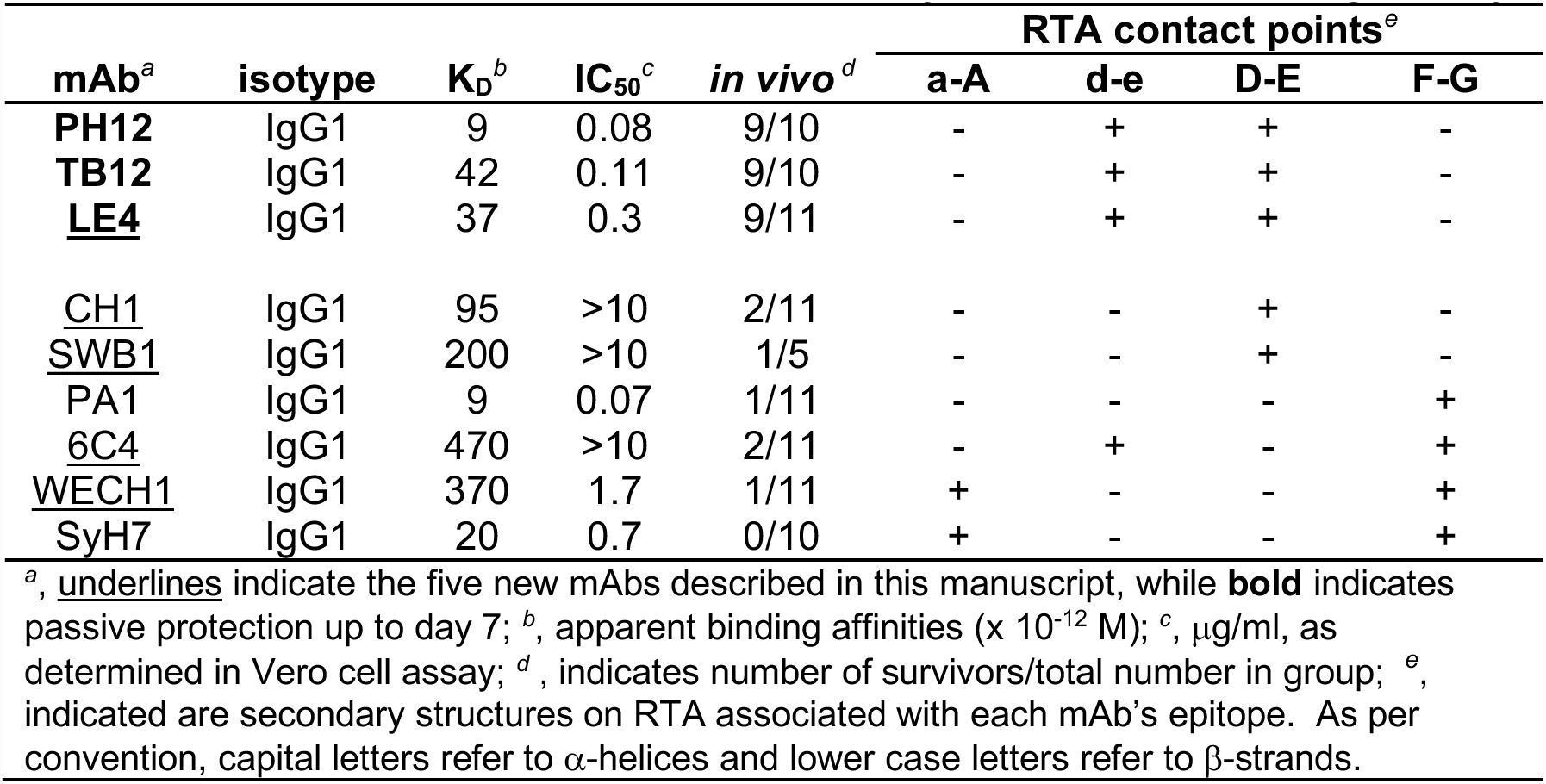
Relationship between epitope specificity and toxin-neutralizing activity

To formally assign the five new mAbs to cluster II, they were subject to a cross competition capture ELISA with a panel of antibodies representing clusters I (PB10, WECB2, R70), II (SyH7, PA1, PH12, TB12), III (IB2), and IV (GD12, JD4). For the most part, the competition results were entirely consistent with the five new mAbs grouping exclusively within cluster II (**Figure 1; Figure S1**). For example, the ability of 6C4 to capture soluble ricin was inhibited by the cluster II mAbs (SyH7, PA1, PH12, TB12), as well as LE4, CH1, WECH1 and SWB1, but not by the representative cluster I, III, or IV mAbs. LE4’s competition profile was also unambiguous except for one instance of non-reciprocal competition with WECB2, one of the three cluster I mAbs. Specifically, soluble WECB2 prevented ricin capture by plate bound LE4, although soluble LE4 did not prevent ricin capture by WECB2 (13). Another anomaly was the non-reciprocal competition between soluble SyH7/SWB1 and plate bound CH1. Nonetheless, the overall competition profiles for WECH1, LE4, CH1, SWB1, and 6C4 are consistent with their grouping within cluster II.

The five new cluster II-specific mAbs were next examined for relative binding affinities (KD) and toxin-neutralizing activities (TNA). As expected, all five mAbs bound ricin toxin by direct ELISA (**Figure 2**). EC_50_ values, determined by capture ELISA using biotin-labeled ricin, ranged from ~4-200 ng/ml (data not shown), while apparent binding affinities, as determined by SPR, ranged from 37-470 pM (**Table 1; Figure S2**). TNA, as determined in a Vero cell cytotoxicity assay ranged from strong for LE4 and WECH1 (IC_50_, 0.3-1.7 μg/ml) to weak (IC_50_, >10 μg/ml) for CH1, SWB1, and 6C4 (**Table 1; Figure 2B**). The four previously described cluster II “legacy” mAbs (PH12, TB12, PA1, SyH7) each had strong TNA.

**Figure 2.**
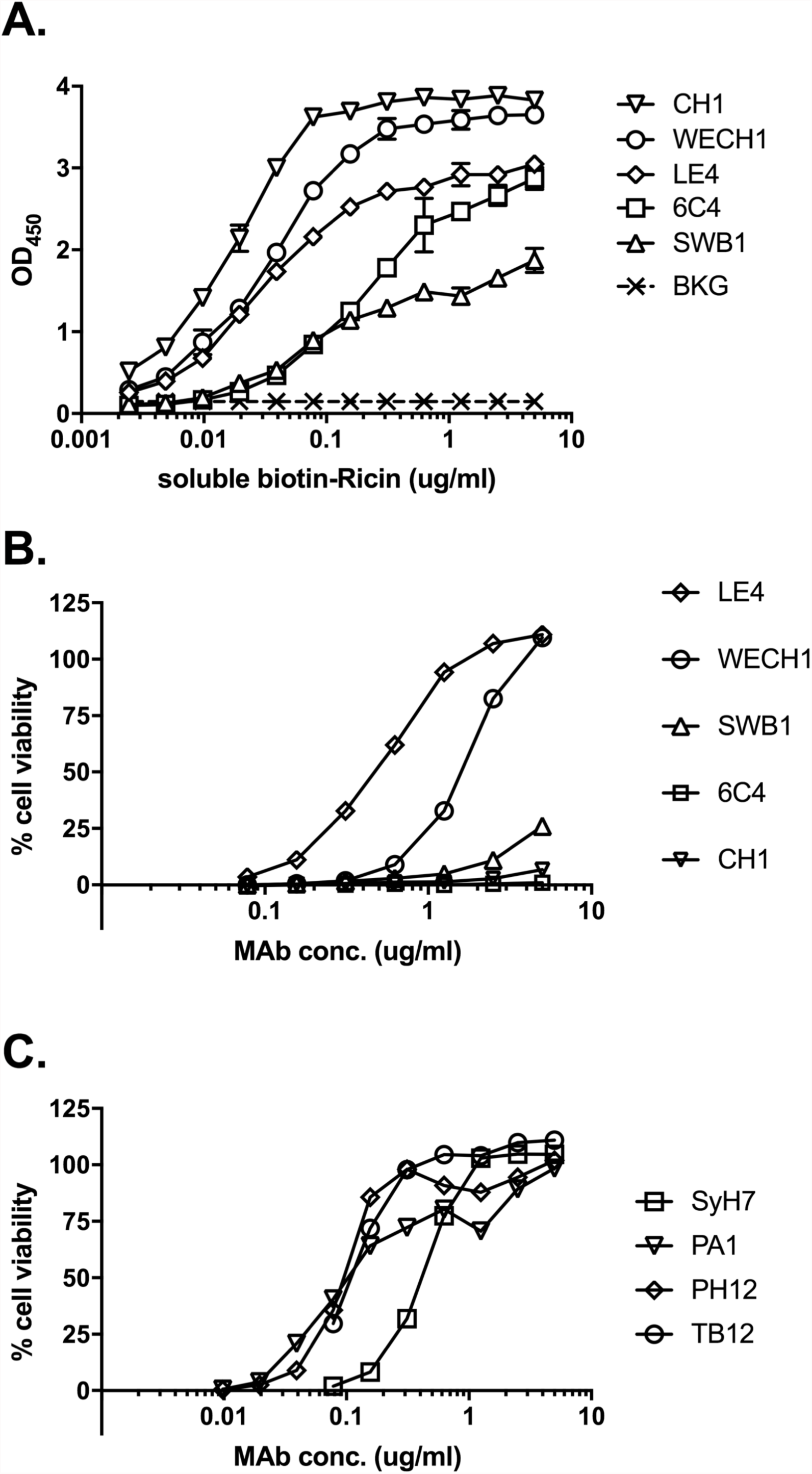
Relative binding profiles and toxin-neutralizing activity of cluster II-specific mAbs. (A) Microtiter plates were coated with indicated mAbs (1µg/ml) and then assessed for the ability to capture biotin-ricin at concentrations on the x-ordinate. Captured biotin-ricin was detected with saturating amounts of avidin-HRP. BKG, background. (B-C) Indicated mAbs at concentrations shown on the X-axis were mixed with ricin toxin (10 ng/ml) and applied to Vero cells for 2 h. The cells were washed and incubated for ~48 h before being assessed for viability. Shown are representative cytotoxicity assays. Actual IC_50_ values are presented in Table 1.

To assess the *in vivo* toxin-neutralizing activities of the five new cluster II mAbs, we performed passive protection studies where groups of mice received individual mAbs by intraperitoneal injection ~6 hr prior to a 10 × LD_50_ ricin challenge by the same route. Mice were monitored for a period of seven days for mortality and weight loss (17, 18). For the sake of comparison, the four cluster II “legacy” mAbs: SyH7, PA1, TB12, and PH12, were also included in the study. It is important to note that previous passive protection studies with the cluster II legacy mAbs were terminated after 3-5 days, not the seven days used here (11, 12).

The results of the passive immunization studies indicated that eight of the nine cluster II mAbs conferred some benefit against ricin intoxication (the exception being 6C4), as compared to control mice that received ricin only (**Table 1,2**; **Figure S3**). However, using survival on day seven as the singular metric, the mAbs stratified into two categories: 1) mice treated with PH12, TB12, or LE4 that were nearly completely protected (90% survival) from ricin-induced death and were by all accounts normal (*e.g.*, feeding behavior, weight gain, grooming) during a several week observation period after the formal completion of the study; and 2) mice treated with one of the other six mAbs (CH1, SWB1, PA1, 6C4, WECH1, SyH7, PA1) that succumbed to ricin intoxication (0-20% survival) by day seven and exhibited significant morbidity during the experimental observation period.

**Table 2.**
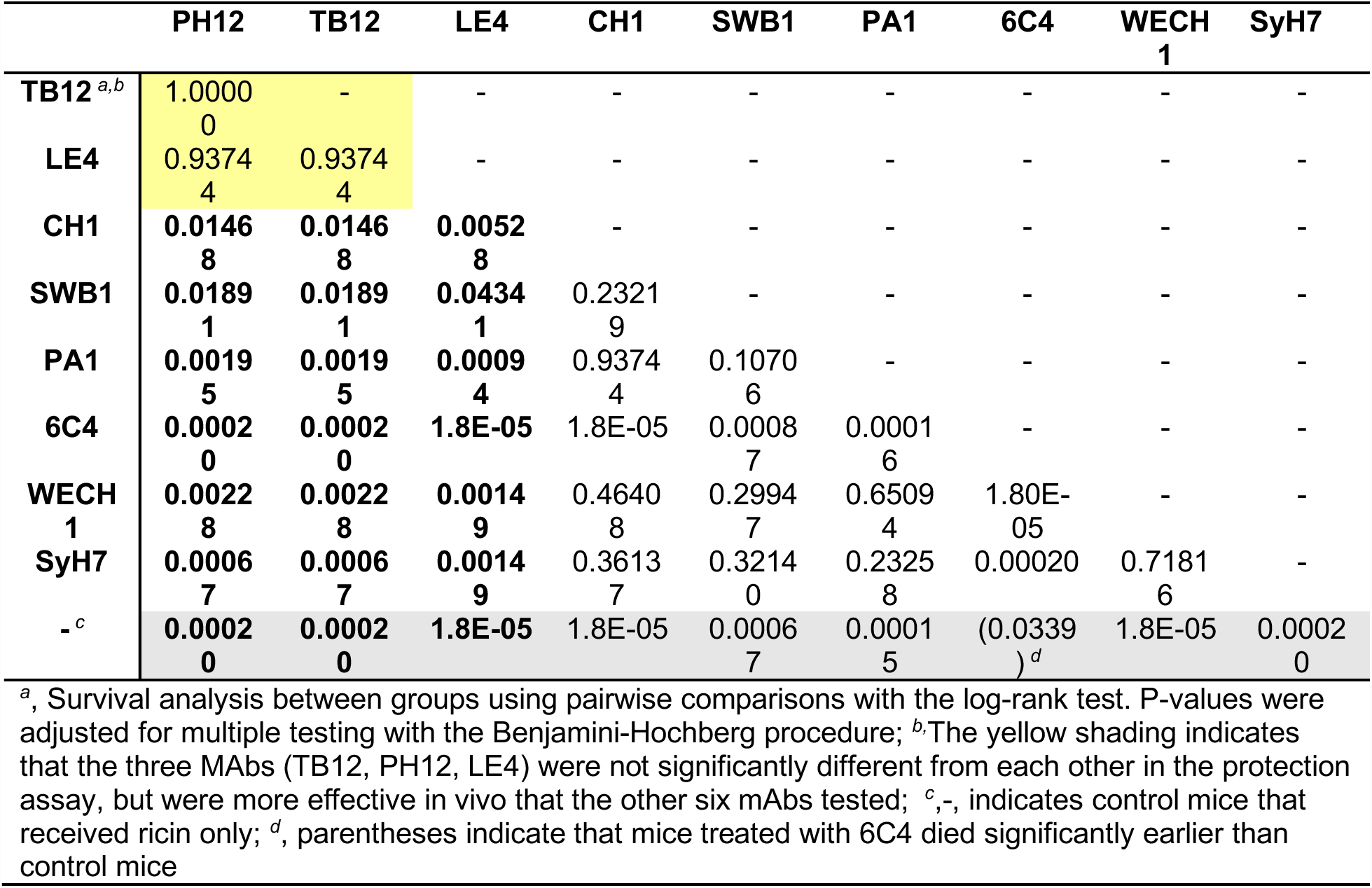
Pairwise analysis of MAbs in passive protection studies

|  | PH12 | TB12 | LE4 | CH1 | SWB1 | PA1 | 6C4 | WECH<br>1 | SyH7 |
| --- | --- | --- | --- | --- | --- | --- | --- | --- | --- |
| **TB12** <sup>a,b</sup> | 1.0000<br>0 | - | - | - | - | - | - | - | - |
| **LE4** | 0.9374<br>4 | 0.9374<br>4 | - | - | - | - | - | - | - |
| **CH1** | 0.0146<br>8 | 0.0146<br>8 | 0.0052<br>8 | - | - | - | - | - | - |
| **SWB1** | 0.0189<br>1 | 0.0189<br>1 | 0.0434<br>1 | 0.2321<br>9 | - | - | - | - | - |
| **PA1** | 0.0019<br>5 | 0.0019<br>5 | 0.0009<br>4 | 0.9374<br>4 | 0.1070<br>6 | - | - | - | - |
| **6C4** | 0.0002<br>0 | 0.0002<br>0 | 1.8E-05 | 1.8E-05 | 0.0008<br>7 | 0.0001<br>6 | - | - | - |
| **WECH**<br>**1** | 0.0022<br>8 | 0.0022<br>8 | 0.0014<br>9 | 0.4640<br>8 | 0.2994<br>7 | 0.6509<br>4 | 1.80E-05 | - | - |
| **SyH7** | 0.0006<br>7 | 0.0006<br>7 | 0.0014<br>9 | 0.3613<br>7 | 0.3214<br>0 | 0.2325<br>8 | 0.00020 | 0.7181<br>6 | - |
| **-** <sup>c</sup> | 0.0002<br>0 | 0.0002<br>0 | 1.8E-05 | 1.8E-05 | 0.0006<br>7 | 0.0001<br>5 | (0.0339<br>)<sup>d</sup> | 1.8E-05 | 0.0002<br>0 |

<sup>a</sup>, Survival analysis between groups using pairwise comparisons with the log-rank test. P-values were adjusted for multiple testing with the Benjamini-Hochberg procedure; <sup>b</sup>The yellow shading indicates that the three MAbs (TB12, PH12, LE4) were not significantly different from each other in the protection assay, but were more effective in vivo than the other six mAbs tested; <sup>c</sup>-, indicates control mice that received ricin only; <sup>d</sup>, parentheses indicate that mice treated with 6C4 died significantly earlier than control mice

The poor outcome of mice treated with CH1, SWB1, and 6C4 was not unexpected considering these mAbs’ relatively modest (or lack of) *in vitro* TNA and sub-optimal binding affinities. However, the failure of SyH7, PA1, and WECH1 to passively protect mice through day seven post-challenge was surprising, considering that their *in vitro* profiles were similar to PH12 and TB12. For example, all four mAbs have similar ricin toxin binding affinities (KD) and roughly equivalent TNA. Thus, it was not immediately apparent what distinguished PH12, TB12, and LE4 from PA1, SyH7, and WECH1.

We had originally assumed, based on competition ELISAs, that SyH7, PA1, PH12, and TB12 recognize the same or nearly the same epitopes (11). This turned out not to be the case, as revealed through recent high-resolution epitope mapping studies using HX-MS. By HX-MS, SyH7 protected RTA’s α-helix A (residues 14-24) and α-helices F-G (residues 184-207). PA1 interacted only with α-helices F-G (residues 184-207), while PH12 and TB12 protected a loop between α-helices D-E (154-164) and a loop between RTA’s β-strands d-e (residues 62-69) (13). The fact that SyH7/PA1 and PH12/TB12 recognize spatially distinct epitopes within cluster II prompted us to define the actual binding sites of the five new cluster II mAbs. Therefore LE4, CH1, SWB1, 6C4, and WECH1 were subjected to epitope mapping by HX-MS (13). A depiction of each 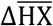 RTA peptide map generated in the absence or presence of mAbs LE4, CH1, SWB1, 6C4, and WECH1 is shown in **Figure 3** and summarized in **Table 3**, with the spatial location of the strongly protected peptides mapped on the surface of RTA shown in **Figure 4**.

**Figure 3.**
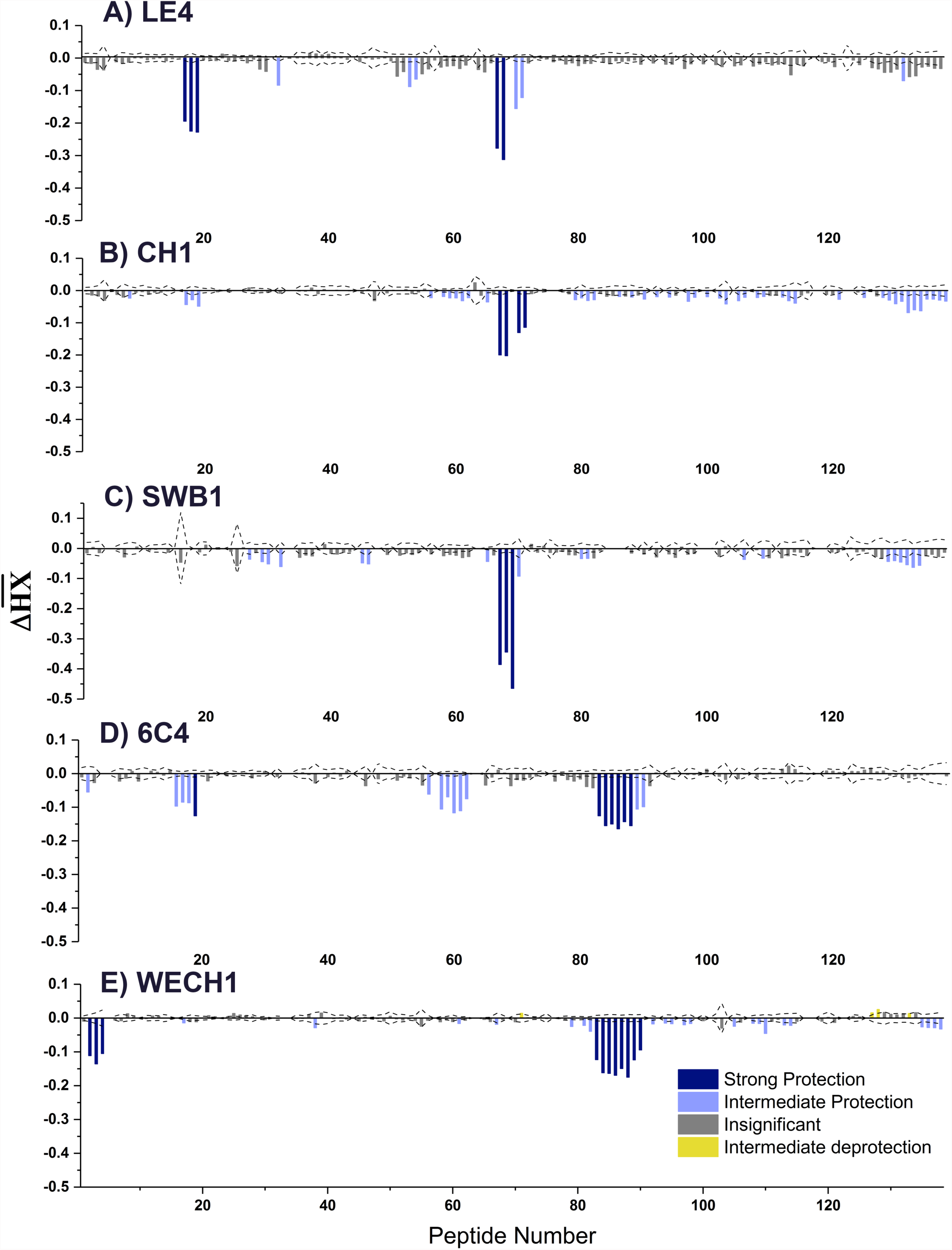
HX-MS analysis of cluster II-specific mAbs LE4, CH1, SWB1, 6C4, and WECH1. Relative levels of protection of RTA peptides by mAbs, as defined by HX-MS (13). The 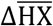 values are clustered using k-means clustering into four categories as follow: strong protection (deep blue); intermediate protection (light blue); insignificant protection (gray); and intermediate deprotection (yellow). The dotted lines represent the threshold for statistically significant changes in hydrogen exchange. The RTA peptides are indexed sequentially from the N-terminus to C-terminus as shown in **Table S1**.

**Figure 4.**
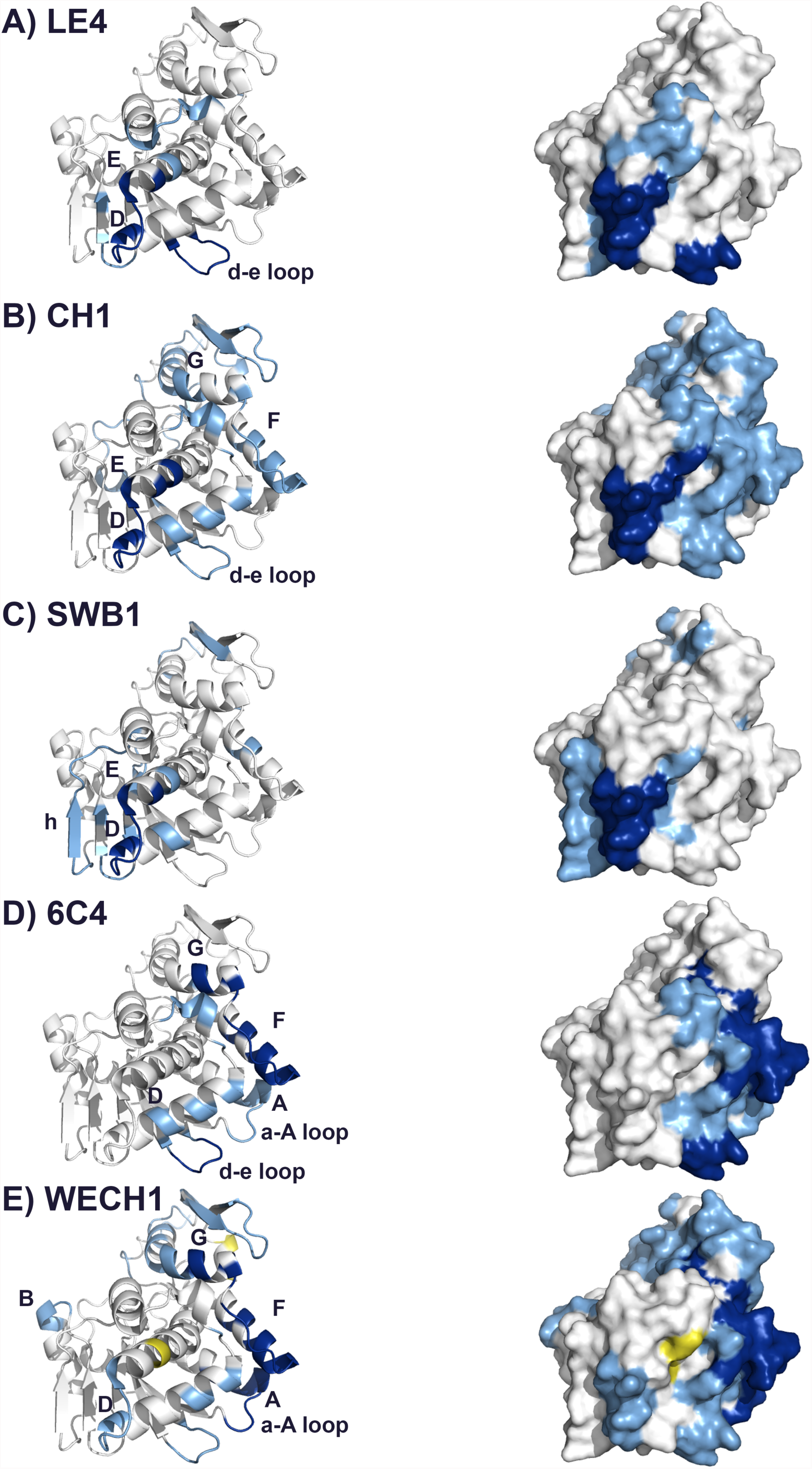
Epitope positioning of cluster II-specific mAbs on RTA. Epitopes on RTA as determined by HX-MS shown using ribbon (left) and surface (right) representations for the five new cluster II mAbs: LE4, CH1, SWB1, 6C4 and WECH1. Degrees of protection are color coded: deep blue, strong protection; light blue, intermediate protection; yellow, intermediate deprotection. Gray indicates insignificant protection. RTA was modeled with PyMol using PDB ID 3SRP.

**Table 3.** HX-MS analysis of cluster II-specific mAbs LE4, CH1, SWB1, 6C4 and WECH1

| mAb | Cluster | Strongly Protected Elements <sup>a</sup> |  |  |
| --- | --- | --- | --- | --- |
|  |  | Peptides | Residues | Proximity |
| LE4 | II | 17-19 | 63-69 | β-strand d |
|  |  | 67,68 | 154-161,163 | α-helices D-E |
| CH1 | II | 67,68,70,71 | 154-161,163,166,167 | α-helices D-E |
| SWB1 | II | 67-69 | 154-161,163 | α-helices D-E |
| 6C4 | II | 19 | 64-68 | β-strand d |
|  |  | 83-88 | 184-187,189-199,210,203,205-206 | α-helices F-G |
| WECH1 | II | 2-4 | 14-20,22-24 | α-helix A |
|  |  | 83-90 | 185-187,189-199,201, 203, 205-207 | α-helices F-G |

<sup>a</sup>, actual exchange profiles and entire RTA peptide list and corresponding residues are provided in **Figure 3**.

HX-MS analysis revealed four distinct protection profiles within the cluster II mAbs examined. LE4 protected RTA peptides encompassing amino acid residues 63-69, which corresponds to a bend between β-strands d and e, and residues 154-161 and 163, which spans the terminus of α-helix D to the proximal residues of α-helix-E. In this respect, LE4 is nearly identical to PH12 and TB12. CH1 and SWB1 were similar to each other in that they protected RTA residues 154-167, corresponding to the loop between α-helix D and α-helix-E. 6C4 strongly protected peptides corresponding to RTA residues 64-68 and 184-206, corresponding to the loop between β-strands d and e (residues 64-68) and the region between α-helices F-G (residues 184-206). Finally, HX-MS analysis indicated that WECH1 contacted RTA residues 14-24, corresponding to a loop between β-strand a and α-helix A and residues 184-207, corresponding to the region between α-helices F-G, a profile identical to SyH7.

A clear pattern emerged when the HX-MS epitope profiles of the five new cluster II mAbs were aligned with the epitopes of the four previously described cluster II mAbs (PH12, TB12, PA1, SyH7), as shown in **Table 1**. Specifically, PH12, TB12, and LE4, the three mAbs with the most potent *in vivo* TNA, had identical HX-MS epitope profiles involving contact with RTA residues 63-69 and 154-163. The other six mAbs either did not engage with residues 63-69 or 154-163, or engaged with just one (but not both) of those particular secondary elements. For example, CH1 and SWB1 protected residues 154-163 but not residues 63-69, while 6C4 engaged with residues 63-69 but not 154-163. SyH7, PA1, and WECH1 do not interact with either residues 154-163 or residues 63-69. It is tempting to speculate that simultaneous binding of an antibody to residues 63-69 and 154-163 (or at least in close proximity to these residues) is a determining factor in antibody potency *in vivo*. We will touch on this topic in the Discussion.

At the level of resolution afforded by HX-MS, PH12, TB12 and LE4 appeared to have the same epitope (13). However, we reported that PH12 and TB12 have different profiles in competition ELISAs with a panel of ~60 VHHs, suggesting that the two mAbs do have distinct binding sites on RTA or different angles of approach (16). To better resolve the LE4 epitope vis a vis PH12 and TB12, we subjected LE4 to similar competition ELISAs with a subset of VHHs. As shown in **Figure 5**, LE4, PH12 and TB12 each had unique competition profiles. For example, PH12 and TB12 did not compete with VHH JNM-E4, whereas LE4 did. TB12 did not compete with VHH JIY-D9, while PH12 and LE4 did. Finally, LE4 and PH12 did not compete with VHHs JIZ-B7 and V5E1, but TB12 did. Thus, PH12, TB12 and LE4 are in fact distinct mAbs with subtle differences in epitope specificity and/or different angles of approach on ricin toxin.

**Figure 5.**
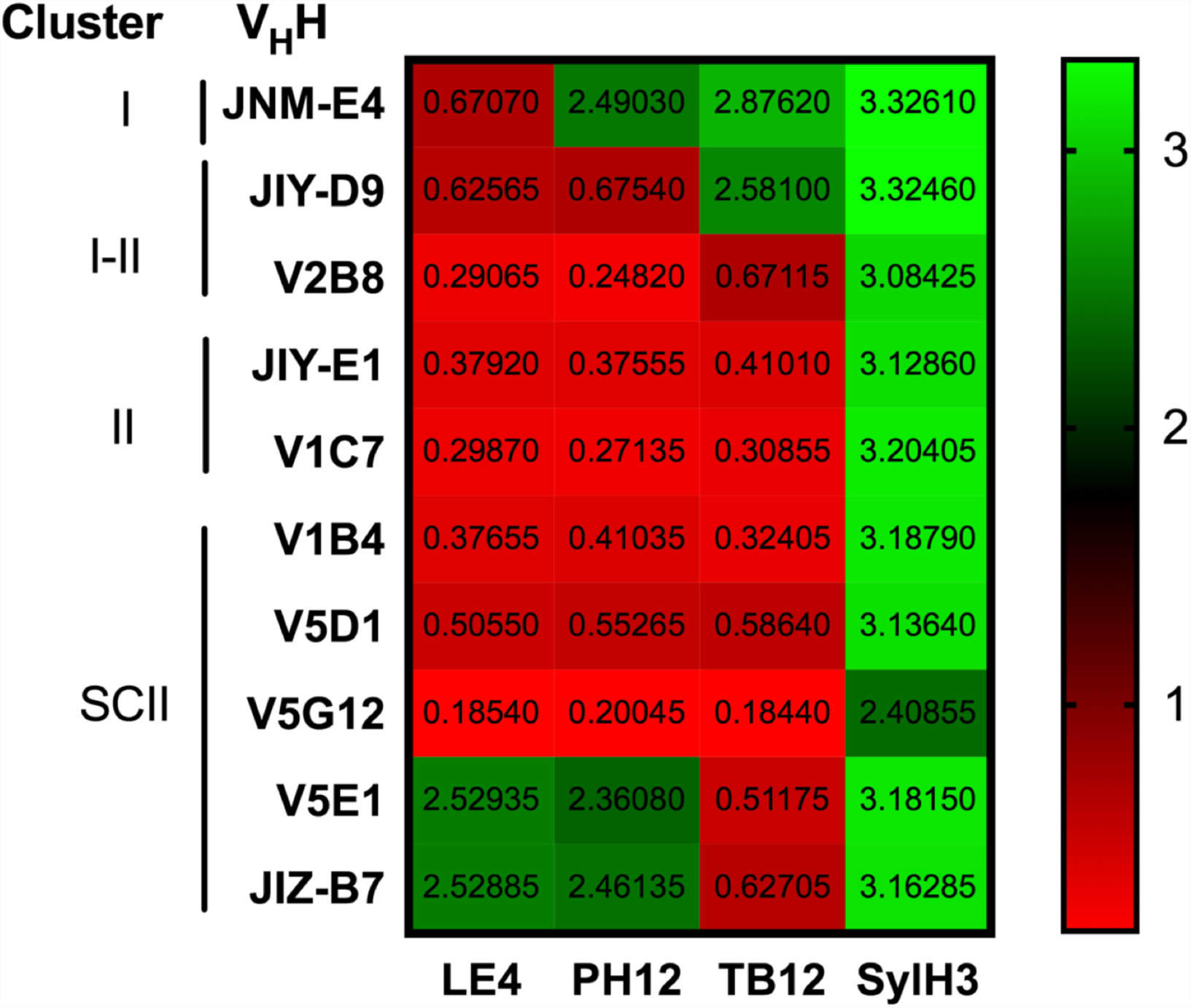
Refinement of cluster II epitopes by VHH competition ELISA. Ricin was captured onto microtiter wells coated with LE4, PH12, or TB12 and then probed with VHHs against epitope clusters I, I/II, II or supercluster II (16). In the far-right column, ricin was captured with SylH3, an RTB-specific mAb that does not to interfere with the panel of VHHs shown here. The heat map is color coded from green (no competition) to red (competition). The actual optical densities obtained from the ELISA as presented in each box. The results shown are from one experiment of three biological replicates with essentially identical results.

## Discussion

In previous reports we delineated four neutralizing hotspots on the surface of ricin toxin’s enzymatic subunit, RTA, that we refer to as epitope clusters I-IV. Cluster I has been interrogated in detail; it is now established that potent toxin-neutralizing activity is associated with antibodies that contact residues within RTA’s α-helix B (97-107) and to a much lesser degree β-strand h (14, 15, 19-21). By contrast, much less is known about cluster II. Recently, we employed HX-MS analysis to localize the epitopes recognized by four so-called cluster II legacy mAbs: TB12, PH12, SyH7, and PA1 (13). In the current report, we have now characterized five additional cluster II mAbs. The five new mAbs, LE4, CH1, SWB1, 6C4, and WECH1, were tested side-by-side with the four legacy mAbs in a mouse model of ricin toxin challenge that was longer in duration than previously studies from our lab. To our surprise, the challenge studies indicated that the cluster II mAbs segregated into two groups (**Table 1**): those that were able to passively protect mice for the duration of the study (LE4, TB12, and PH12), and 2) those that were not (CH1, SWB1, PA1, 6C4, WECH1, and SyH7). The two groups varied in their relative binding affinities and in their capacity to neutralize ricin *in vitro*, However, the most interesting difference between these groups was their contact sites on RTA. The protective mAbs, LE4, TB12, and PH12 each make strong contact with RTA residues 63-69 and 154-163, while the other six mAbs did not.

We conclude that cluster II as a whole constitutes a relatively large patch situated on the back side of RTA, relative to the active site (**Figure 6A**). When ricin holotoxin is positioned with RTB at its base, the TB12, PH12, and LE4 epitopes are located towards the toxin’s apex along with CH1 and SWB1, while the other epitopes within cluster II are situated in closer proximity to RTB. Unfortunately, there are no landmarks within or in proximity to cluster II that might explain why antibody occupancy in this region differentially attenuates ricin’s toxicity. In other words, why do TB12, PH12, and LE4 have potent *in vivo* toxin-neutralizing activity, while PA1 and SyH7 do not? TB12, PH12, and LE4 make two primary contact points with RTA: residues 63-69 and residues 154-163. Residues 63-69 correspond to a bend between β-strands d and e, which are part of a six stranded β-sheet (strands a-h) that dominant the first of RTA’s three folding domains (10). No particular function has been ascribed to this six stranded b-sheet, even though it clearly is integral to the overall tertiary structure of RTA. Elimination of the bend between β-strands d and e by site-directed mutagenesis (Δ62-66) did not impact RTA’s ability to depurinate ribosomes in a cell free assay (22). Residues 154-163 encompass the C-terminus of α-helix D, a short intervening loop (residues 157-161), and the proximal residues of α-helix-E. Removal of residues 152-156 or 157-161 did not impact RTA activity *in vitro*, although perturbing α-helix-E renders the subunit inactive (22). We do postulate that the surface area delineated by cluster II is important for ricin cytotoxicity, possibly playing a role in intracellular transport. We have demonstrated, for example, that SyH7, when bound to ricin, affects the efficiency of toxin transport from the plasma membrane to the TGN (23). SyH7 also interferes with *in vitro* protein disulfide isomerase (PDI)-mediated reduction of the disulfide bond that links RTA to RTB, an event that normally occurs in the ER (24). Unfortunately, neither TB12, PH12, nor LE4 have been tested yet in these types of assays.

**Figure 6.**
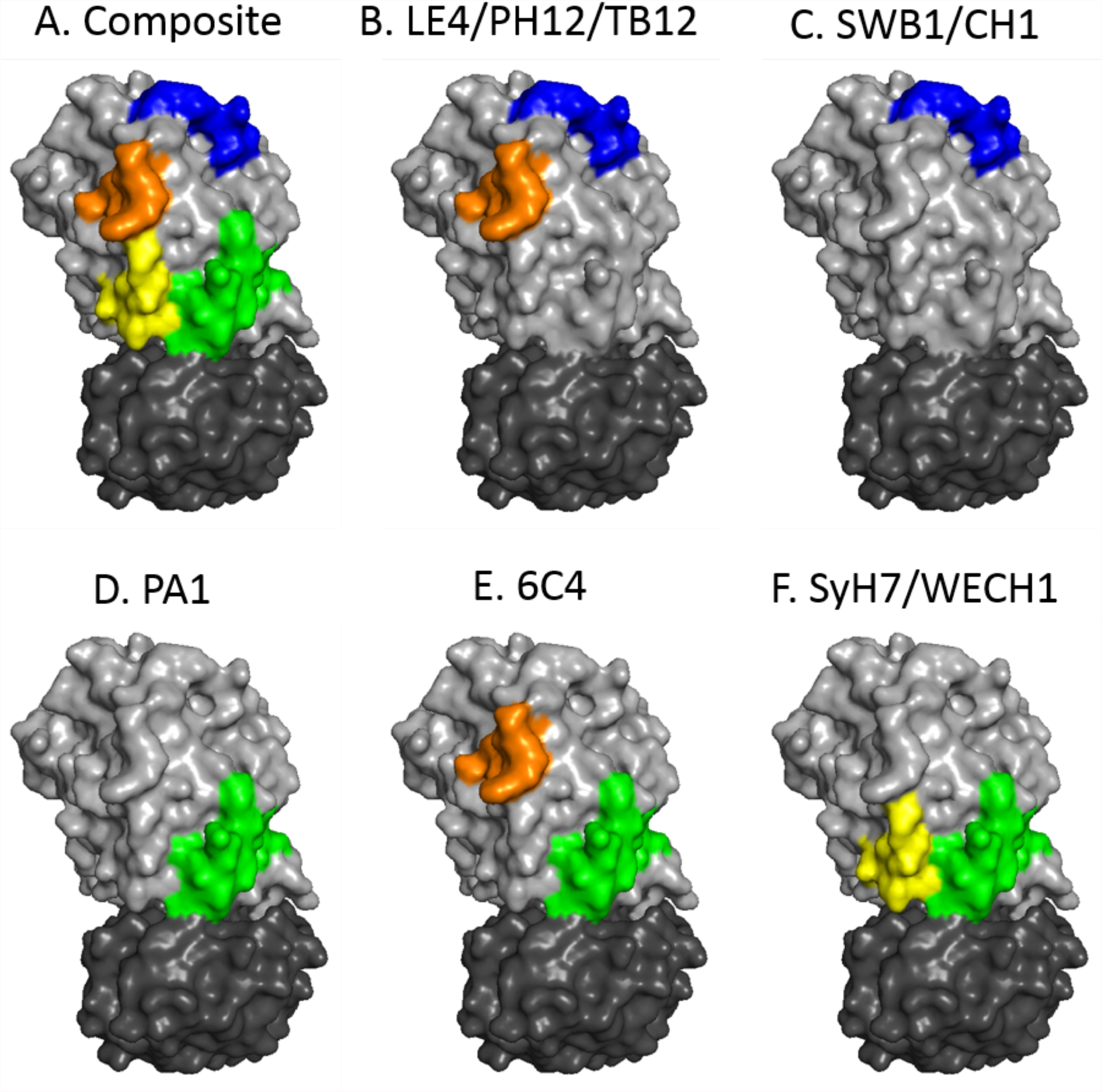
Surface depiction of cluster II epitopes on the surface of ricin toxin. (A) Composite image of the four cluster II structural contact points listed in **Table 1** mapped onto the structure of ricin (PDB ID 2AAI) using PyMol. (B-F) Individual epitopes for indicated mAbs. Colors: RTA, light gray; RTB, dark gray; β-strand a-α-helix A, yellow; d-e, orange; D-E, blue; F-G, green.

The resolution afforded by HX-MS analysis is such that we were unable to distinguish differences in epitope specificity between LE4, PH12 and TB12. However, competition ELISAs with a collection of V_H_Hs demonstrate that LE4, PH12 and TB12 are indeed different from each other. Based on available high resolution V_H_Hs epitope maps, generated in some cases by X-ray crystallography, we can speculate as to how LE4, PH12 and TB12 may differentially engage RTA at residues 63-69 and 154-163. All three mAbs compete with V_H_Hs V1C7 and JIY-E1, which are known from X-ray crystallography to contact RTA’s d-e and D-E loops. By contrast, JNM-E4 competes with LE4, but not PH12 or TB12. JNM-E4 is grouped within epitope cluster I and targets the “top” of RTA. Thus, LE4 likely approaches ricin from the top down, relative to PH12 and TB12. TB12 competes V5E1 and JIZ-B7, whereas LE4 and PH12 do not. V5E1 and JIZ-B7 recognize epitopes at the RTA-RTB interface (25, 26), suggesting TB12 approaches RTA from the side or even underside. Finally, we postulate that PH12 attacks ricin at an angle somewhere between LE4 and TB12, based on competition with JIY-D9. Ultimately, assigning exact epitopes to LE4, PH12 and TB12 will require X-ray crystal structures of the mAbs in complex with RTA or ricin holotoxin.

Based on HX-MS analysis, the other six mAbs in cluster II (CH1, SWB1, PA1, 6C4, WECH1, and SyH7) recognize at least four different epitopes on RTA. Three of the mAbs, PA1, WECH1 and SyH7, have strong *in vitro* TNA, while the other three are devoid of activity, which is explained in large part by differences in relative binding affinities. In previous studies we concluded that SyH7 and PA1 were in fact protective in our mouse model. However, it is now apparent that those conclusions were incorrect because the experiments were terminated prematurely. Moreover, we failed to use body weight as a marker of morbidity, which other investigators have used successfully (18). From the current study it is clear that mice treated with even relatively high doses of SyH7, CH1, SWB1, PA1, 6C4 or WECH1 begin to lose weight almost immediately after ricin challenge, whereas mice treated with TB12, PH12, or LE4 maintained normal body weights. While the actual basis for ricin-induced death following systemic toxin exposure remains unknown, it most likely pertains to liver or kidney failure (27). Therefore, it is interesting that SyH7 is unable to neutralize ricin when mouse liver sinusoidal endothelial cells or Kupffer cells (KCs) are the target cells, at least ex vivo (B. Mooney and N. Mantis, *manuscript in preparation*). We have just started examining the other cluster II mAbs for the ability to protect LSECs and KCs from ricin.

Finally, it should be noted that cluster II is actually more complicated than has been presented up to this point in the Discussion. Recent comprehensive epitope mapping studies along with X-ray crystallography have revealed that certain cluster II antibodies recognize quaternary epitopes involving residues on RTA and RTB, an aggregate of epitopes we refer to as supercluster II (SCII) (16, 25, 26). A prime example is V5E1, a camelid VHH whose CDR1 and CDR2 elements contact RTA along α-helix A (residues 18-32), α-helix F (182-194), and the F-G loop, which explains competitive interference with SyH7 (26). At the same time, V5E1’s CDR3 straddles the RTA-RTB interface and docks in close proximity to RTB’s high affinity Gal/GalNAc lectin element. Conversely, JIZ-B7 is an example of a VHH whose primary target is RTB, but whose binding to ricin holotoxin is inhibited by SyH7 (which defines a cluster II antibody) (16, 25). The X-ray crystal structure of JIZ-B7 bound to ricin holotoxin is not available, although the structures of six other SCII VHHs bound to ricin holotoxin have been solved (M. Rudolph, D. Vance, N. Mantis, *manuscript in preparation*).

## Materials and Methods

### Chemicals, Reagents and Cell lines

Ricin (*Ricinus communis* agglutinin II) was purchased from Vector Laboratories (Burlingame, CA) and dialyzed against PBS using a Slide-A-Lyzer dialysis cassette (Pierce, Rockford IL) prior to use in animal experiments and cytotoxicity assays. Goat serum (New Zealand origin) was purchased from Gibco-Life Technologies (Carlsbad, CA). Cell culture media was prepared by the Wadsworth Center Media Services facility. MAbs were affinity purified from hybridoma supernatants by endotoxin-free protein G chromatography at the Dana Farber Cancer Institute’s Monoclonal Antibody Core facility (Boston, MA). African green monkey kidney (Vero) cells were obtained from the American Type Culture Collection (ATCC, Manassas, VA). All other chemicals were purchased from Sigma-Aldrich, Inc (St. Louis, MO), unless otherwise specified.

### Animal care and B-cell hybridoma production

All mouse experiments were conducted in accordance with the Wadsworth Center’s Institutional Animal Care and Use Committee (IACUC) guidelines. Mice were housed under conventional, specific pathogen-free conditions. Six-week-old female BALB/c or Swiss Webster mice (Taconic Biosciences; Albany, NY) were administered sub-lethal amounts of ricin by intraperitoneal (i.p.) injection as follows: 0.1 µg on days 0, 10 and 20; 0.2 µg for BALB/c and 0.3 µg for SW on day 35. Mice were retro-orbitally bled on day 45, serum was tested by ELISA and toxin-neutralization assay to confirm seroconversion. As a final boost, mice were injected i.p. with the equivalent of ~10 × LD_50_ ricin (2 µg) and then euthanized four days later by CO_2_ asphyxiation. Splenocytes were fused with mouse myeloma cells using HybriMax polyethylene glycol (PEG). Fusion products were seeded into 96-well tissue-culture treated plates and cultured/selected in RPMI (Gibco) media supplemented with UltraCruz hybridoma cloning supplement (Santa Cruz Biotechnology; Dallas, TX) containing fetal calf serum, oxaloacetate, sodium pyruvate, bovine insulin, HAT (hypoxanthine-aminopterin-thymidine) and penicillin-streptomycin. HAT was gradually replaced with HT (hypoxanthine-thymidine), after which surviving hybridomas secreting antibodies of interest were cloned by limiting dilution and expanded in RPMI media without HT. Bulk hybridoma line expansions were cultured in serum-, protein-, and antibiotic-free CD media (Gibco) and resulting supernatants were cleared by filtration before being submitted for affinity purification.

### Direct and competitive ELISAs

ELISAs were performed as described (28). For direct ELISAs, NUNC Maxisorb F96 microtiter plates (Thermo Scientific, Pittsburgh, PA) were coated with 1 µg/ml MAb or ricin diluted in PBS, pH 7.4. Plates were blocked with 2% goat serum (Gibco) in PBS-tween (0.1%). Medium containing MAb or biotinylated-ricin was then applied to wells neat or diluted into block solution, and incubated at room temperature (RT). Horseradish peroxidase (HRP)-labeled goat anti-mouse IgG-specific polyclonal antibodies (SouthernBiotech; Birmingham, AL) or avidin-HRP (Thermo Scientific) were used as secondary reagents, along with 3,3’,5,5’- tetramethylbenzidine (TMB; Kirkegaard & Perry Labs, Gaithersburg, MD) as colorimetric detection substrate; a 1M phosphoric acid solution was added to each well to stop the reaction. Plates were read on a VersaMax spectrophotometer and analyzed using Softmax Pro 5.4.5 software (Molecular Devices, Sunnyvale, CA). EC_50_ values were determined by nonlinear regression of soluble ricin binding curves using least squares method within the ECanything function of GraphPad Prism 7.01.

Epitope profiling immune-competition capture (EPICC) was performed as follows: Immulon 4HBX 96-well microtiter plates (Thermo Scientific) were coated with “capture” mAb (1 μg/ml) diluted in PBS, pH 7.4 and incubated at room temperature. Wells were then blocked with 2% goat serum/PBS-tween (0.1%) solution overnight at 4°C. The biotinylated-ricin (biotin-R) limiting concentration used for this capture assay was equal to the EC90 concentration for each coated mAb (range= 30-200 ng/ml); this concentration was kept constant across all wells and each biotin-R solution was diluted in blocking solution containing 2% goat serum. Tenfold excess of “competitor” mAb solutions were made in separate tubes: mAbs were diluted to 10 μg/ml in their respective EC_90_ values in solution, incubated 15 min, then applied to wells in duplicate. A series of at least 4 wells per coated mAb were overlaid with biotin-R EC_90_-only solution as 100% binding controls for the purpose of calculating binding inhibition. Plates were incubated at room temp. for 1 h. Wells were then washed 3 with PBS with 0.1% Tween-20 and overlaid with HRP-avidin (1 ug/ml) followed by TMB. Plates were analyzed with a VersaMax spectrophotometer, using Softmax Pro 5.2.5 software. Ricin binding inhibition was calculated as a percentage of BR binding to the capture mAb, where: [100-(OD_450_*C*/OD_450_*B*)*100]= *% ricin binding inhibited by competitor*; *C*, competed, *B*, biotin-R EC90 control.

### Vero cell toxin neutralization assay

Ricin TNA were performed as described (28). Opaque tissue culture treated 96-well plates (Corning) containing confluent layers of Vero cells were treated with ricin (10 ng/ml), ricin-MAb mixtures (in duplicate), or medium alone (as negative control), then incubated 2 h at 37°C. Initial treatments were then aspirated; wells were overlaid with DMEM supplemented with 10% fetal bovine serum (FBS) and penicillin-streptomycin and incubated for 48 h at 37°C. Cell viability was assessed using CellTiter-GLO reagent (Promega; Madison, WI); plates were read on a SpectraMax L luminometer (Molecular Diagnostics) and analyzed using SoftMax Pro 5.2.5 software. 100% viability was defined as the average value of all wells treated with medium only. IC_50_ values were determined by nonlinear regression of cell viability curves using least squares method within the ECanything function of GraphPad Prism 7.01.

### Surface plasmon resonance (SPR)

MAb association and dissociation rates for ricin toxin were determined by SPR using the ProteOn XPR36 (Bio-Rad Inc.; Hercules, CA), as described (21). For ricin immobilization, general layer compact (GLC) chips were equilibrated in running buffer PBS-0.005% Tween (PBS-T, pH 7.4) at a flow rate of 30 µl/min. Following EDAC (200 mM) sulfo-NHS (50 mM) activation (3 min), ricin was diluted in 10 mM sodium acetate (pH 5.0) at two different concentrations (4 µg/ml and 2 µg/ml) and immobilized (2 min). A third vertical channel received only acetate buffer and served as a reference channel. The surfaces were deactivated using 1 M ethanolamine (5 min). The ProteOn MCM was then rotated to the horizontal orientation for antibody experiments. Each mAb was serially diluted in running buffer and injected at 50µl/min for 180s, followed by 1 to 3 hr of dissociation. After each experiment, the chip surface was regenerated with 10 mM glycine, pH 1.5 each at 100 µl/min for 18 s, until the RU values returned to baseline. All kinetic experiments were performed at 25°C. Kinetic constants for the antibody/ricin interactions were obtained with the ProteON Manager software 3.1.0 (Bio-Rad Inc.).

### Passive Protection Studies

Monoclonal antibodies (10 µg or 25 µg) were diluted in endotoxin-free PBS and administered by i.p. injection in a final volume of 0.4 ml to 8-week-old female BALB/c mice (Taconic Biosciences, Albany, NY). Six hours later mice received the equivalent of ~10 × LD_50_ of ricin (2 µg per mouse) by i.p. injection. Following ricin challenge, mice were weighed and scored for morbidity twice daily for 7 days; mice were euthanized when they became overtly moribund and/or weight-loss was >20% pre-challenge weight, as mandated by the Wadsworth Center’s IACUC.

### Hydrogen exchange mass spectrometry (HX-MS)

HX-MS experiments were conducted using a LEAP H/DX PAL system (Carrboro, NC) and a quadrupole time of flight (QTOF) mass spectrometer (Agilent, Santa Clara, CA). HX-MS workflow and data processing for mapping epitopes of recombinant RTA by mAbs were carried as described (13). For reasons of safety, HX-MS was conducted on a recombinant version of RTA carrying two attenuating point mutations (Y80A, V76M), known as RiVax (29). For simplicity, we will simply refer to it a recombinant RTA in this study. In brief, regions of RTA that exhibited significantly slower (protection) or faster (deprotection) HX in the presence of mAbs were identified using a combination of k-means clustering and significance testing based on time-averaged HX measurements, 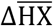, quantifying the difference between the mAb-RiVax complex and unbound recombinant RTA. Unlike in the previous work, here the results are filtered for solvent accessibility of the RTA residues (Toth et al., *manuscript in preparation*). k-means clustering was used to classify the effect of the mAb on the HX of RTA* from strongly protected to deprotected. Strongly protected regions were used to define the epitopes.

### Statistical analysis

Differences in survival between groups were determined with Kaplan-Meier analysis and log-rank testing. Pair-wise comparisons between groups were performed with log-rank tests and the resulting p-values were adjusted for multiple comparisons with the Benjamini-Hochberg procedure to control the false discovery rate. For all analyses p-values of <0.05 were considered significant. All statistical analysis was carried out in GraphPrad Prism 7 (GraphPad Software, San Diego, CA), or R version 3.4.2 (30).

## Acknowledgements

We thank Dr. Renjie Song in the Wadsworth Center’s Immunology Core facility for assisting with SPR studies. We gratefully acknowledge Dr. Jenny Tang of the Wadsworth Center’s Cell Culture Facility and Dr. Ed Greenfield of the DFCI Monoclonal Antibody core facility for assistance in purification of mAbs. We also thank the Wadsworth Center animal care staff for pre- and post-study animal maintenance.

## Footnotes

Research reported in this work was supported by Contract No. HHSN272201400021C from the National Institutes of Allergy and Infectious Diseases, National Institutes of Health. The content is solely the responsibility of the authors and does not necessarily represent the official views of the National Institutes of Health. The funders had no role in study design, data collection and analysis, decision to publish, or preparation of the manuscript.

